# USP30 inhibition improves mitochondrial health through both PINK1-dependent and independent mechanisms

**DOI:** 10.1101/2025.02.03.636341

**Authors:** Matthew G. Williamson, Rachel Heon-Roberts, Sarah N. J. Franks, Elliot Mock, Hannah B. L. Jones, Elena Britti, Ana Malpartida, Mahmoud Bassal, Martha Lavelle, Jack Connor, Aina Mogas Barcons, Katherine Hammond, Katrina Savory, Pavendeep Rai, Anna Lavayssiere, William McGuinness, Natalie Sepke, Remya Raghavan-Nair, Jane Vowles, Iolanda Vendrell, Franziska Guenther, Benedikt M. Kessler, Sally Cowley, Katherine England, Emma Murphy, John Davis, Richard Wade-Martins, Brent J Ryan

## Abstract

Mitochondrial dysfunction is a key feature of many pathologies, including Parkinson’s disease. The selective vulnerability of dopaminergic neurons is thought to be influenced by mitochondrial dysfunction and mutations in the mitophagy regulating proteins PINK1 and Parkin that are known to cause early-onset Parkinsonism in an autosomal recessive manner.

Augmentation of mitophagy through inhibition of USP30 may be a viable therapeutic strategy for a number of diseases including Parkinson’s. USP30 inhibition has been demonstrated to augment PINK1/PRKN mitophagy but also potentiate basal mitophagy to support the removal of dysfunctional mitochondria. Therefore, long-term de-regulation of mitophagy has been proposed to lead to mitochondrial depletion.

We have used an integrated approach across cell lines, primary neurons and iPSC-derived dopaminergic neuronal cultures to assess the short and long-term effects of USP30 inhibition on mitochondrial health and neuronal activity. We investigated the dependence of USP30 inhibition phenotypes on the PINK1/Parkin pathway using genetic ablation and in iPSC-derived neurons from Parkinson’s patients with PINK1 or PRKN mutations.

Loss of USP30 through CRISPR/Cas9-mediated knockout resulted in increased basal and depolarisation-induced mitophagy in SH-SY5Y cells. Loss of USP30 or pharmacological inhibition altered mitochondrial morphology and led to increases in membrane potential and ATP levels with decreased oxygen consumption, suggesting that USP30 loss results in a more efficient mitochondrial network. These changes in morphology were found to be independent of PINK1 or Parkin.

Chronic pharmacological inhibition of USP30 or CRISPRi-mediated knockdown of USP30 did not impact dopaminergic neuronal activity, as assessed by electrophysiological profiling, but did potentiate depolarisation-induced mitophagy in primary and iPSC-derived neuronal cultures. We observed minimal changes in mitophagy levels in iPSC-derived dopaminergic neurons from Parkinson’s patients with PINK1 or PRKN mutations that were independent of the ability to produce p65Ub. Importantly, within this experimental paradigm, pharmacological USP30 inhibition increased depolarisation-induced mitophagy in both PINK1 and PRKN patients to the same extent as control neurons.

These results support a role for USP30 in modulating the trigger threshold for mitophagy and suggest that USP30 inhibitors may be beneficial in patients with impairments in PINK1/Parkin-mediated mitophagy.

## Introduction

Parkinson’s disease (PD) is the fastest-growing neurodegenerative disorder worldwide, affecting 2% of individuals over 60 years old [1]. The disease is characterized by the preferential loss of dopaminergic neurons in the substantia nigra pars compacta (SNpc). The cellular pathology underpinning the loss of dopaminergic neurons is multifactorial and includes both proteostasis and mitochondrial dysfunction. Mitochondrial dysfunction is linked to monogenic and idiopathic forms of Parkinson’s as well through toxins, which may be driven by dopaminergic neuron physiology [2]. In particular, homozygous or compound heterozygous mutations in the genes *PINK1* and *PRKN* (Parkin) are associated with early-onset Parkinsonism [3–5]. PINK1 and Parkin cooperate in a feed-forward mechanism to build ubiquitin chains on mitochondria in response to mitochondrial depolarisation to promote mitophagy [6, 7]. Pathogenic mutations in PINK1 or PRKN have been demonstrated to impair depolarisation-induced mitophagy [8]. Therefore, strategies to upregulate mitophagy have been pursued for a number of diseases including Parkinson’s.

USP30 is a mitochondrially-tethered deubiquitylase that removes ubiquitin from mitochondrial outer-membrane substrates in a spatially-restricted manner, restraining PINK1/Parkin-dependent mitophagy [9, 10]. The structural and mechanistic basis of USP30 activity has been elucidated in exquisite detail in a range of models. Additionally, USP30 has also been demonstrated be present on the surface of peroxisomes, to regulate their turnover (pexophagy) [11, 12].

Knockdown or knockout of USP30 has been demonstrated to have a number of effects on cellular biology and mitochondrial health including increasing both basal and depolarisation-induced mitophagy [13, #1096]. However, in iNeurons, Ordureau *et al*. found decreased levels of basal mitophagy, as assessed using mKeima-XL [10], suggesting that the effects of USP30 inhibition may be cell-type dependent. Recent studies have demonstrated that USP30^-/-^ mice are viable and resistant to motor deficits and decreased dopamine levels induced by adenoviral delivery of A53T-SNCA [14].

Several small-molecule covalent and non-covalent USP30 inhibitors have been reported across a range of cellular and animal models [13–17]. Many of these compounds have demonstrated the ability to increase depolarisation-induced mitophagy through PINK1/PRKN and rescue disease models such a mouse AAV-A53T-SNCA model [14] and isogenic PRKN KO lines [17].

The selectivity profile of a small-molecule benzosulfonamide-containing, non-covalent USP30 inhibitor (*(S)*-CMPD-39) has been profiled in neuroblastoma cells demonstrating selectivity for USP30 against a panel of 40 deubiquitylases [16, 18]. The compound has been demonstrated to induce increased ubiquitylation of USP30 substrates TOM20 and SYNJ2BP as well as enhance mitophagy and pexophagy in cell lines [16].

In the present study we further investigated the effects of knockout, knockdown and pharmacological inhibition of USP30 in neuroblastoma cell lines, primary rat neuronal cultures and iPSC-derived dopaminergic neuronal cultures. Using these models, we investigated if USP30 inhibition caused deleterious effects on mitochondrial and neuronal function and the dependence of phenotypes caused by loss of USP30 activity on the PINK1/Parkin pathway to understand how USP30 inhibitors may be beneficial in Parkinson’s patients with PINK1 or PRKN mutations.

## Methods

### Generation of stable knockout SH-SY5Y lines using CRISPR/Cas9

CRISPR/Cas9 mediated knockout of USP30, PINK1 and PRKN was achieved in inducible Cas9 expressing SH-SY5Y using lentiviral expression of sgRNA as previously described [19]. sgRNA sequence (GTTCACCTCCCAGTACTCCA) was used to cut in Exon 3 of USP30 achieving a knockout score of 87% (polyclonal line). Single clones were picked from this population and USP30 levels characterized by sequencing and western blotting to establish homozygous KO (USP30^-/-^) and heterozygous KO (USP30^+/-^) lines (Figure S1C). Three separate sgRNA targeting Exon 1 of PINK1 (TCTTTCTGGCCTTCGGGCTA, CCTCATCGAGGAAAAACAGG, CATCGAGGAAAAACAGGCGG) were multiplexed achieving a polyclonal knockout score of 75%. A single sgRNA targeting Exon 6 of PRKN (GATCGCAACAAATAGTCGGA) was used to achieve a polyclonal knockout score of 34%. From these pools, single clones were isolated by limiting dilution and resulting monoclonal lines were characterized by sequencing and western blotting to establish homozygous KO (USP30^-/-^, PINK1^-/-^and PRKN^-/-^) and heterozygous KO (USP30^+/-^) lines (Figure S1C & S2A-B).

### Measurement of mitophagy and mitochondrial morphology using mitoQC

pLVX mCherry-GFP-mtFIS1(101-152) (mitoQC) was a gift from Prof Ian Ganley (MRC PPU, Dundee, UK). SH-SY5Y were generated by lentiviral infection followed by FACS of mitoQC expressing cells generating a polyclonal stable line. Primary cortical cultures were transduced with Mito-QC reporter lentivirus at a MOI of 2 immediately before plating on DIV 0 with the media was changed 24 h later.

To analyze mitochondrial morphology, nuclei were stained with NucBlue (R37605, Invitrogen) and mitoQC positive cells were identified using Harmony analysis software (Revvity). In presence of a BFP reporter, nuclei of mitoQC positive cells were stained with HSC NuclearMask deep Red 1X for 30 min (ThermoFisher). Briefly, mitochondria morphology was assessed using SER ridge filter of GFP positive channel after removal of mCherry positive puncta, morphology measurements for area, roundness, width, length and width/length ratio, were calculated. Mitophagy events were calculated by counting mCherry spots.

Mitochondrial morphology was determined at the single mitochondrion level by extracting mitochondrial morphology parameters on a single object basis (Harmony software). Data were processed using R Studio.

### Measurement of mitophagy using mKeima

mKeima induction was controlled using a tetracycline-inducible promoter (TRE3G) as previously described [20]. Stable SH-SY5Y cell line was engineered using lentiviral infection of cells with mKeima viral particles at MOI of 2, followed by FACS sorting of positive cells following 1h doxycycline induction to generate a polyclonal cell population. Briefly, mKeima expression was induced in SH-SY5Y or iPSC-DaN with 1.5µg/ml for 24h prior to imaging. mKeima was imaged using an Opera Phenix CLS (Revvity). In SH-SY5Y, mitophagy was quantified by the number of ‘red’ (ex. 561 nm) spots per cell. In iPSC-DaN, in which there is less clear contrast between the ‘green’ (ex. 421 nm) and ‘red’ (ex. 561 nm) signal the mitophagy index was quantified using the ratio of mt-mKeima total red area (ex. 561 nm)/mt-mKeima total green area (ex. 421 nm), emission of keima was measured at 570-630 nm.

### Generation of rat cortical primary cultures

Primary cortical cultures were generated as previously described [21]. Briefly, a minimum of three Sprague–Dawley rats (P2–P5) were pooled for each experiment, in accordance with UK Home Office regulations, under the Animals (Scientific Procedures) Act of 1986. Brains were extracted, meninges were removed single cells were obtained through trypsin digest. Cells were counted using an automated cell counter NucleoCounter NC-250 (ChemoMetec, Denmark). Cells were seeded at 5 × 10^5^ and kept in the incubator at 37°C, 5% CO_2_, 95% relative humidity and 50% of medium was replaced every 2 days. All assays were performed on DIV 14–15 cells.

### Participant recruitment

Participants gave signed informed consent to mutation screening and derivation of iPSC lines from skin biopsies (Ethics committee: National Health Service, Health Research Authority, NRES Committee South Central, Berkshire, UK, REC 10/H0505/71). PINK1 and PRKN mutation lines were reprogramed from previously described lines Table 1 [22–29].

**Table 1.** Description of iPSC lines used in this study.

| Line name | Sex | Age | Disease Status | Mutation in PINK1/PRKN | iPSC line Reference |
| --- | --- | --- | --- | --- | --- |
| SFC-065-03 | M | 65 | Control | - | [28] |
| SFC-156-03 | M | 62 | Control | - | [20] |
| SFC-180-01 | F | 60 | Control | - | [45] |
| SFC-840-03 | F | 67 | Control | - | [20] |
| SFC-841-03 | M | 36 | Control | - | [30] |
| SFC-856-03 | F | 78 | Control | - | [30] |
| SFC-822-03 | F | 31 | Parkinson's | PINK1<br>(Homo Q456X) | This study |
| SFC-823-03 | F | 61 | Parkinson's | PINK1<br>(Homo V170G) | This study |
| SFC-824-03 | M | 38 | Parkinson's | PINK1<br>(Homo Q456X) | This study |
| SFC-825-03 | F | 53 | Parkinson's | PINK1<br>(Homo Q456X) | [29] |
| SFC-826-03 | F | 47 | Parkinson's | PINK1<br>(Homo Q456X) | [29] |
| SFC-817-03 | M | 64 | Parkinson's | PRKN<br>(c.1072Tdel + Ex7del<br>DEL) | [27] |
| SFC-818-03 | M | 15 | Parkinson's | PRKN<br>(R275W + Ex4del) | [27] [29] |
| SFC-821-03 | F | 38 | Parkinson's | PRKN<br>(R275T + C352R) | This study |

### Differentiation of iPSC-derived dopaminergic cultures (iPSC-DaN)

iPSC-DaN were differentiated as previously described [20, 30]. Detailed methodology for the differentiation of iPSC-DaN can be found here (dx.doi.org/10.17504/protocols.io.q26g7y1jqgwz/v1). Briefly, iPSC were differentiated by plated at a density of 1.5 × 10^5^ cells/cm^2^ on Geltrex (Life Technologies) for two days. iPSCs were patterned to ventral midbrain precursors for 10 days then expanded for up to 19 days in KSR and NNB medium (1:3) supplemented with LDN193189 (100 nM; Merck) and CHIR99021 (3 μM, Tocris). Cells were passaged four times 1:2 with Accutase up to four times and plated with 10 μm Y-27632 onto Geltrex-coated plates. Cells were frozen at DIV 10 + 18 in CellBanker 2 (11914, Amsbio). Following the expansion of ventral midbrain precursor cells (DIV 11), media was changed to NB medium containing Neurobasal medium, B27 and 2 mM l-glutamine (Life Technologies) and differentiated into dopaminergic neuronal cultures. Cells were replated at DIV 20 and seeded at 2.5 × 10^5^ cells/cm^2^, followed by 15 days of maturation. All experiments were carried out at DIV 35-40 and data were gathered from two to three independent differentiations.

### CRISPRi knockdown of USP30

Top scoring sgRNA targeting USP30 were identified using data from Horlbeck et al [31] and cloned into lentiviral backbone encoding a nuclear-localised mTagBFP2 and puromycin selection cassette [32]. Knockdown of USP30 in SH-SY5Y was achieved using SH-SY5Y stably expressing dCAS9-BFP-KRAB [33] followed by puromycin selection. Knockdown of USP30 in iPSC-derived neurons was achieved by infection of sgRNA into D20 iPSC-DaN.

### Assessment of neuronal activity in iPSC-DaN

Axion Biosystems Cytoview 48-well MEA plates (16 electrodes/well, cat no: M768-tMEA-48W) were coated two days before cell plating (D18) with poly-L-ornithine (8 µL droplet/ well, Sigma-Aldrich, P4957, 0.01% solution) and incubated overnight at 37 °C. PLO was removed and Biolaminin (8 µL droplet/well, 25 µg/mL diluted in DPBS++, Biolamina, LN521) was added and again incubated overnight at 37 °C. On the day of plating (D20), 30,000 iPSC-derived dopaminergic neurons in 6 µL Neurobasal maturation media with ROCK inhibitor were spotted on the recording electrodes and allowed to settle for 1 hour at 37 °C followed by careful addition of maturation media with ROCK inhibitor (200 µL/well). Optional lentivirus was added in the media at an MOI of 4. After 2 days (D22), cultures were treated with mitomycin C for 1 hour to remove undifferentiated cells. From D24 half media changes (100 µL per well) were carried out three times a week using BrainPhys + B27 maturation media including all growth factors with the addition of laminin (2 µg/mL final concentration, Sigma-Aldrich, L2020). MEA recordings (2 min, 37 °C and 5% CO2, sampling frequency = 12.5 kHz) were acquired at least once a week on a Maestro PRO (Axion Biosystems) using AxIS Navigator v3.6.2.2 from D30 onwards to assess neuronal firing activity. Plates were equilibrated in the Maestro PRO for 10 min prior to recording. Recordings were always performed one day after media changes. An adaptive threshold spike detector set to 5.5 standard deviations was applied to the raw data. RAW files were analysed with AxIS Navigator to produce metric files describing firing and bursting activity. The following parameters were used: Active electrode criterion = 5 spikes/min; Burst detection: inter-spike interval threshold, maximum inter-spike interval = 100 ms, minimum number of spikes = 5. Weighted mean firing rate was calculated by averaging the mean firing rate across only active electrodes. Further analysis of the metric files was performed with AxIS Metric Plotting Tool v2.4.4 and Excel, and testing for statistical significance with Graphpad Prism v8. Treatment with (*S*)-CMPD-39 (3 µM) was performed after recording a baseline at D82 followed by a full feed (100 µL). (*S*)-CMPD-39 was included in half media changes for 21 days. 1 healthy control iPSC line (SFC856-03-04) was used with six replicate wells per condition.

### Cellular oxygen consumption

iPSC-derived DA neuronal cultures were plated in a XF96 Polystyrene Cell Culture Microplate (Seahorse Bioscience) on day 20 and further matured until day 35 before analysis. SH-SY5Y were plated at a density of 2x10^4^ cells/well 24h prior to assaying. On the day of the assay, the assay medium was prepared fresh using the XF Base Medium (Agilent Technologies) supplemented with 10 mM glucose (Merck), 1 mM sodium pyruvate (Merck) and 2 mM l-glutamine (Thermo Fisher Scientific). At least 1 h before the assay, the cells were washed once with the assay medium and then incubated at 37°C in a non-CO2 incubator. Three baseline recordings were made, followed by sequential injection of the ATP synthase inhibitor oligomycin (Oligo; Merck, 1 μM), the mitochondrial uncoupler p-trifluoromethoxyphenylhydrazone (FCCP; Merck, 1 μM), the Complex I and III inhibitors Rotenone and Antimycin A (R/A; Merck, 5 μM) and the glucose analog 2-deoxyglucose (2-DG; Merck, 50 mM). Mitochondrial respiration and glycolytic activity of DA or SH-SY5Y cultures were measured using a Seahorse XFe96 Analyzer (Agilent Technologies) and normalized to total well protein content (Pierce BCA protein assay kit).

### Mitochondrial membrane potential

To detect changes in mitochondrial membrane potential (ΔΨm), cells were incubated with 5 μM JC-10 (AAT Bioquest, Inc.) at 37°C for 1 h. The fluorescence intensities were measured using the multi-mode plate reader PHERAstar FSX (BMG Labtech) by fluorescence. Monomer was measured at ex: 514 nm em: 529 nm and aggregated form at ex 585 em 590 nm. Membrane potential was defined as ΔΨm before/after addition of 10 μM carbonyl cyanide 3-chlorophenylhydrazone (CCCP; Merck).

### ATP production measurement

Cellular ATP levels were assayed using CellTiter-Glo 3D Cell Viability Assay (Promega) in 96-well plates. iPSC-DaN were assayed at 3X10^4^ cells/well in half-area 96-well plates (Greiner). SH-SY5Y were assayed at a density of 5x10^4^ cells/well. The substrate was added to cell culture medium and 100 µl was transferred to a 96-white plate and incubated for 25 min at room temperature, shielded from light. To measure ATP production, luminescence was measured using the multi-mode plate reader PHERAstar FSX (BMG Labtech) and measurements were normalized to cell number. Cell number was measured prior to the start of the assay by adding Hoechst nuclear marker (NucBlue Live ReadyProbes Reagent, R37605; Invitrogen) to cells and imaging live (Opera Phenix High Content Screening System; Revvity).

### Mitochondrial reactive oxygen species

Mitochondrial reactive oxygen species (ROS) was assessed using MitoSOX probe (M36008, Life Technologies). Cells were incubated in half-area 96-well plates with 2.5 μM MitoSOX Red Mitochondrial Superoxide Indicator and nuclear marker Hoechst (NucBlue Live ReadyProbes Reagent, R37605; Invitrogen) for 10 min at 37°C with 5% CO_2_. Cells were washed with media and immediately MitoSox fluorescence was imaged at ex: 488 em: 650–760 nm, using an Opera Phenix High Content Screening System (Revvity).

### Assessment of p65Ub levels by HTRF

P65Ub levels were assessed in cell lysates using HTRF Human & Mouse Phospho-Ubiquitin (Ser65) Detection Kit (64UBIS65PEG, Revvity)

### Immunoblotting

Levels of proteins were determined in cell lysates by western blotting as previously described [19]. Briefly, cell pellets were lysed in RIPA buffer and subjected to SDS-PAGE before transfer onto PVDF membranes using a TransBlot Turbo (Bio-Rad). Membranes were probed overnight with primary antibodies as follows: mouse anti-β-Actin-HRP (Abcam #49900), p65Ub antibody (ABS1513, Millipore).

### Activity based probe assays

HA-Ub-PA synthesis was carried out as previously described [34, 35]. (S)-CMPD-39/DMSO was incubated with SH-SY5Y cells at the indicated concentrations/incubation times. Cells were washed 3 x with PBS, collected at 200 x g for 10 minutes and lysed in 50 mM Tris Base, 5 mM MgCl2.6 H2O, 0.5 mM EDTA, 250 mM Sucrose, 1 mM DTT by vortexing with acid washed beads (1:1 V/V) 10 times (30 seconds vortexing, 1-minute break on ice). Lysates were clarified at 600 g for 10 minutes at 4 °C and protein concentration was determined by BCA. Lysates were incubated with a saturating concentration of HA-Ub-PA for 5 minutes at 37 °C. Reactions were then quenched with the addition of Laemmli buffer and subject to western blotting. USP30: Atlas antibody HPA016952, GAPDH: Invitrogen MA5-15738, Anti-HA: Roche 11666606001. USP30 activity was extracted from western blots using densitometric quantification in image studio lite (version 5.2.5). HA-Ub-PA labelled USP30 was taken as a percentage of the sum of USP30 bands +/− HA-UB-PA to establish 100 % activity, and reduced activity upon inhibition.

## Results

### Genetic ablation of USP30 potentiates mitophagy resulting in improved mitochondrial function

To assess the effects of USP30 on mitochondrial health, in cells with endogenous Parkin expression, we used CRISPR-Cas9 to generate monoclonal heterozygous and homozygous USP30 KO SH-SY5Y lines (USP30^-/+^ and USP30^-/-^) (Supplemental figure 1A-C).

To assess the effect of USP30 knockout on mitophagy initiation, we assessed levels of the PINK1 substrate p65Ub, using a high-content imaging assay. Knockout of USP30 resulted in a gene-dosage-dependent potentiation of p65Ub formation upon CCCP-induced mitochondrial depolarization (Figure 1A-B). However, basal p65Ub levels were unaltered in USP30 KO SH-SY5Y lines (Supplemental 1D).

**Figure 1.**
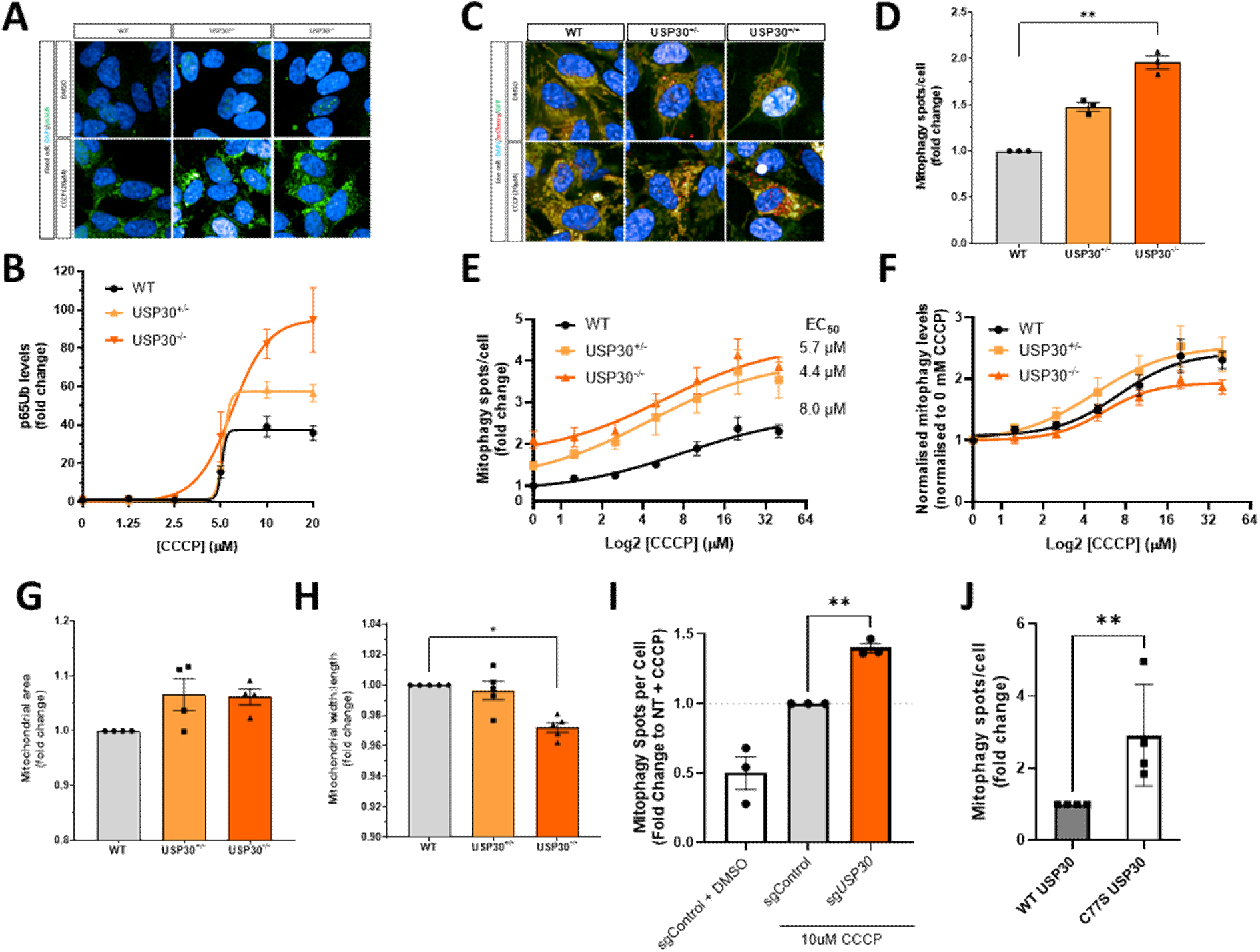
Loss of USP30 induces basal mitophagy and enhances depolarisation-induced mitophagy dependent on USP30 catalytic activity. (A-B) p65Ub levels in USP30^-/-^ cells. Cells were treated for 6h with CCCP and p65Ub levels measured by immunofluorescence A) Representative images B) quantification of p65Ub levels (n=3). C-G) quantification of mitophagy and mitochondrial morphology using mitoQC in USP30^-/-^ cells under basal and CCCP-stimulated conditions. C) Representative images D-E) quantification of mitophagy as assessed by red (mitolysosome) spots under D) basal (unstimulated) or E) CCCP-stimulated conditions (n=3) F) mitophagy rates (from panel E) in CCCP-stimulated cells normalised within line to DMSO. Quantification of G) mitochondrial network area and H) Mitochondrial morphology, as assessed by mitochondrial width:length ratio (n=5). I) Mitophagy assessed using mtKeima in dCas9-BFP-KRAB cells expressing non-targeting sgRNA (sgControl) or sgRNA targeting USP30 (sgUSP30) stimulated with CCCP for 6h (n=3). J) Quantification of mitophagy using MitoQC in USP30^-/-^ cells expressing WT or C77S USP30 (n=4) and Samples were run in triplicate. *p<0.05, **p<0.005, ***p<0.001, one-way ANOVA followed by Dunnett’s multiple comparison to WT, error bars represent mean +/- SEM.

Using the fluorescence reporter mitoQC, mitophagy and mitochondrial morphology were assessed under basal conditions and after mitochondrial depolarisation with the uncoupler CCCP in cells with homo/heterozygous knockout of USP30. Loss of USP30 resulted in a two-fold increase in basal mitophagy and an increase in depolarization-induced mitophagy (P<0.001 vs WT cells ANOVA with Tukey post-test) (Figure 1C-E). Normalization of mitophagy levels within lines demonstrates that mitophagy levels with increasing concentrations of CCCP may be driven by increased basal mitophagy rates rather than increased responses to depolarisation (Figure 1F). Together, these data support previous reports [11], which demonstrate that the increase in stimulated mitophagy observed in USP30^-/-^ cells was driven by initial substrate ubiquitination rather than acceleration of PINK1/Parkin-mediated mitophagy.

To address the hypothesis that increased basal mitophagy drives mitochondrial depletion in conditions of USP30 loss, we assessed if mitochondrial number, morphology and function was impaired in USP30 KO lines. In contrast with the hypothesis of mitochondrial depletion, the area of mitochondria not associated with lysosomes was increased in USP30^-/-^ cells concomitant with a decrease in the mitochondrial width:length ratio (Figure 1 G-H), indicating a longer, less fragmented mitochondrial network.

Induction of mitophagy and morphology changes were recapitulated in polyclonal USP30-/- lines in which we demonstrated increased mitophagy and p65Ub induction (S1E-I). In addition, CRISPRi knockdown of USP30 levels resulted in increased CCCP-induced mitophagy relative to non-targeting sgRNA (sgControl) as assessed by the fluorescent reporter mtKeima P<0.01, paired t-test) (Figure 1I).

To confirm the dependence of these phenotypes on the deubiquitylase activity of USP30, we reconstituted USP30-/- cells with WT USP30 or catalytically-inactive (C77S) USP30 constructs expressing a BFP reporter. Reconstitution of USP30-/- cells with WT USP30 decreased mitophagy and increased mitochondrial fragmentation relative to cells expressing catalytically-inactive (C77S) USP30 (Figure 1J & S1J-M).

Given the increase in mitochondrial area and decrease in fragmentation, the effect of USP30 knockout on mitochondrial function was assessed. Loss of USP30 resulted in an increase in mitochondrial membrane potential and cellular ATP levels in a gene-dosage-dependent manner (Figure 2 A-B). Basal mitochondrial ROS levels were unchanged in USP30 KO cells, with a modest decrease in rotenone-stimulated ROS production (Figure 2C-D). Basal mitochondrial oxygen consumption rate was decreased in USP30^-/-^ cells (Figure 2 E-F). Consistent with these changes being dependent on USP30 catalytic activity, reconstitution of USP30-/- cells with WT USP30 expression raised mitochondrial oxygen consumption rates relative to C77S USP30 (Figure 2 G-H).

**Figure 2.**
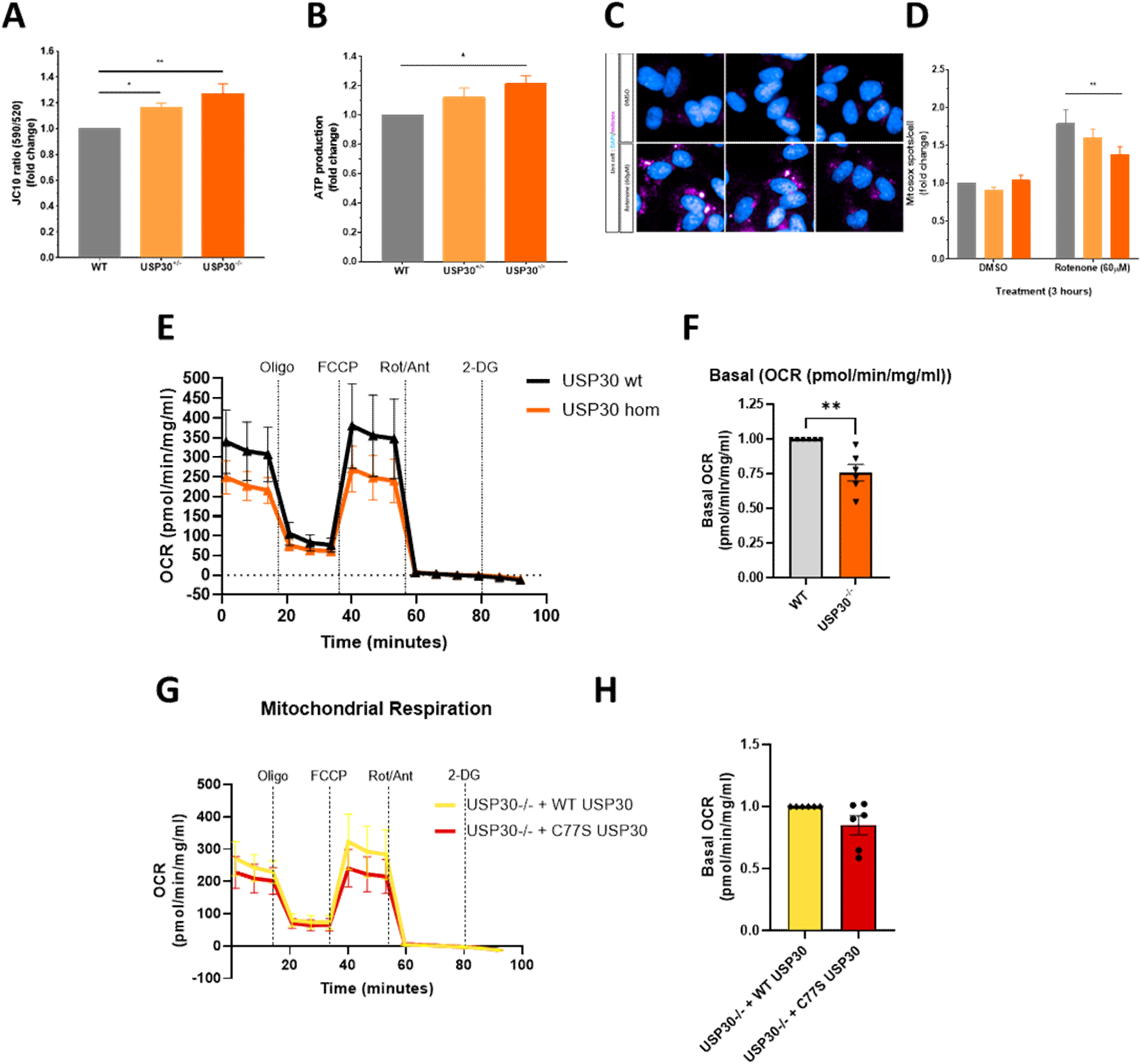
Loss of USP30 increases mitochondrial efficiency. **A)** Mitochondrial membrane potential was assessed using the potentiometric dye JC-10 (n=4) **B)** Cellular ATP levels assessed using luciferase-based luminescence (n=4) **C-D)** Mitochondrial ROS assessed by MitoSox staining under basal and Rotenone-stimulated conditions (60µM) **C)** Representative images and **D)** quantification of ROS levels (n=3). E-G) Oxygen consumption rates (OCR) in WT and USP30^-/-^ cells assessed using extracellular flux assay (n=6) **E)** normalised OCR after injection of mitochondrial modulators **F)** quantification of basal OCR. **G-H)** OCR in USP30^-/-^ cells expressing WT or C77S USP30 (n=4). Error bars represent mean +/- SEM

Together, these data suggest that whilst USP30 knockout increases basal mitophagy, this does not result in a depletion of mitochondria but in a more efficient mitochondrial network in SH-SY5Y.

### Pharmacological USP30 inhibition recapitulates the effects of loss of USP30

To understand if these phenotypes can be recapitulated with pharmacological USP30 inhibition, the non-covalent USP30 inhibitor compound-39 (CMPD-39) was used. *(S)*-CMPD-39 (the *(S)*-enantiomer) is exquisitely potent against USP30, however, the corresponding *(R)*-enantiomer is >1000-fold weaker against USP30 and was used as an inactive control (Figure 3 A). *(S)*-CMPD-39 effectively inhibited USP30 activity in a dose- and time-dependent manner for up to 24h (Figure 3B-C). *(S)*-CMPD-39 potentiated CCCP-induced p65Ub generation in a dose-dependent manner, with enantiomer (R)-CMPD-39 showing markedly decreased potency (EC50∼0.2 µM and 2µM, respectively) (Figure 3 D-E). However, *(S)*-CMPD-39 did not further potentiate p65Ub in USP30-/-cells indicating that *(S)*-CMPD-39 potentiates p65Ub through USP30 (Figure 3 F). Consistent with USP30 KO and p65Ub potentiation, *(S)*-CMPD-39 potentiated CCCP-induced mitophagy to a greater extent than (R)-CMPD-39 (Figure 3G).

**Figure 3.**
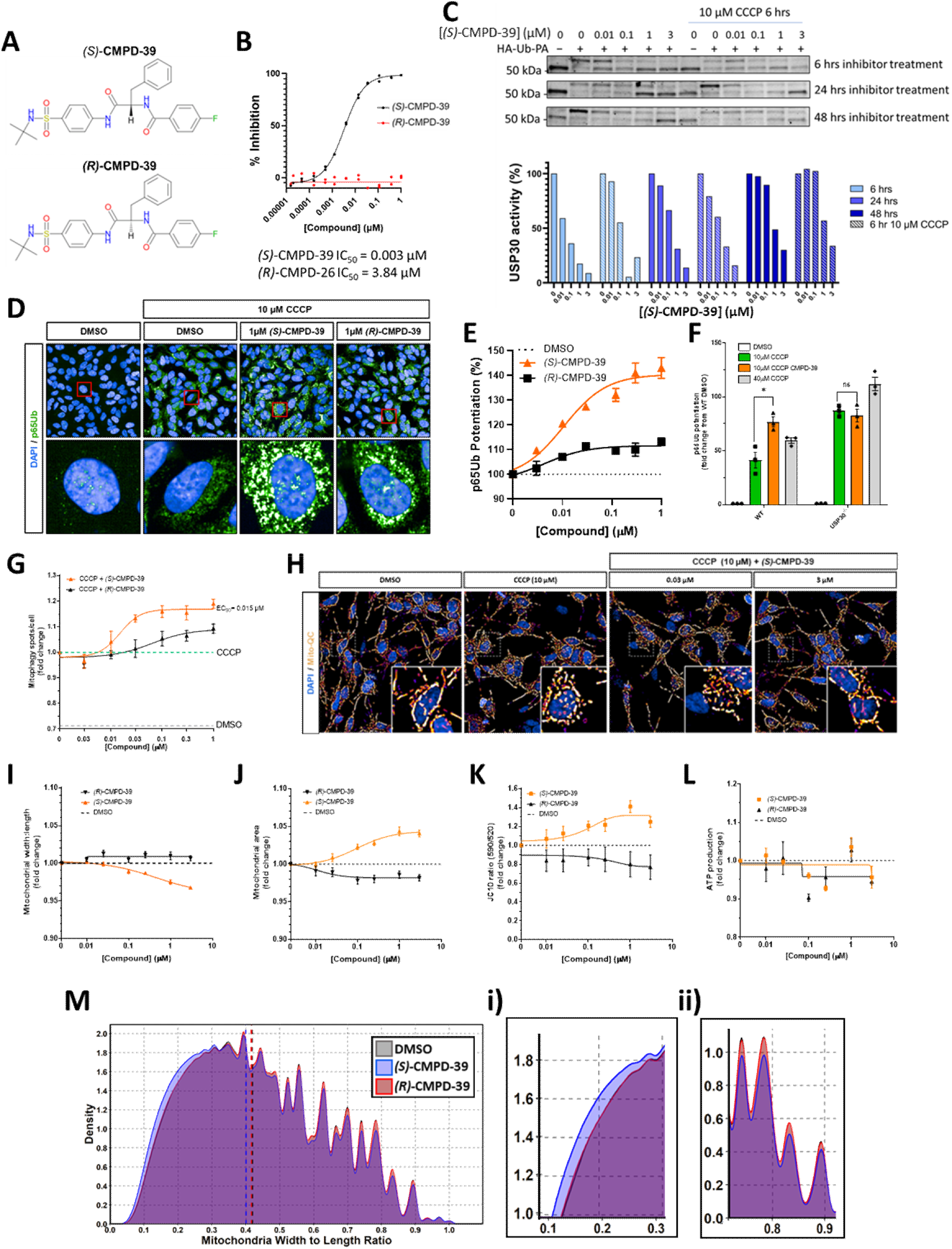
Pharmacological inhibition of USP30 improves mitochondrial health via distinct mechanisms. **A)** Structures of *(S)*-CMPD-39 and its inactive stereoisomer *(R)*-CMPD-39 **B)** Dose-dependent inhibition of USP30 by *(S)*-CMPD-39 and *(R)*-CMPD-39. **C)** time dependence of USP30 inhibition in SH-SY5Y determined using a HA-Ub-PA probe and assessed by western blotting. **D-E)** p65Ub levels in untreated (grey dashed line) and CCCP-treated cells (green dashed line) treated with increasing concentrations of active and inactive USP30 inhibitors 6h treatment. **F)** p65Ub in WT or USP30-/- cells treated with 3µM *(S)*-CMPD-39 in the presence/absence of CCCP stimulation. **G-J)** Mitophagy and mitochondrial morphology as assessed using mitoQC **G)** representative images of mitochondrial network morphology identified using SER ridge parameter in DMSO and CCCP-treated cells **H-J)** quantification of **H)** mitophagy (n=4). **I)** mitochondrial morphology (width:length ratio) and **J)** mitochondrial area, as quantified using non-lysosomal MitoQC signal in cells treated with active/inactive USP30 inhibitors for 48h. **K)** mitochondrial membrane potential assessed using JC-10 fluorescence in cells treated for 48h with active/inactive USP30 inhibitors (n=3). **L)** Cellular ATP levels assessed using luciferase-based assay in cells treated for 48h with active/inactive USP30 inhibitors **M)** Histogram showing mitochondrial morphology quantified on a single mitochondrial level in SH-SY5Y treated with active/inactive USP30 inhibitors. Zoom of **i)** elongated and **ii)** fragmented mitochondria.

In line with observations in USP30-/-, pharmacological USP30 inhibition resulted in a dose-dependent decrease in mean mitochondrial width:length, indicating that mitochondria are on average longer after USP30 inhibitor treatment (Figure 3 H-I). Similarly, *(S)*-CMPD-39 resulted in a dose-dependent increase in mitochondrial area and increased ψm but were independent of large changes in cellular ATP levels (Figure 3J-L).

To understand the mechanism of the increased mean mitochondrial length (decreased width:length ratio) observed at the cell population level, we investigated if this population effect is driven by increased mitochondrial fusion or decreased numbers of fragmented mitochondria. We assessed the distribution of the mitochondrial morphology at the single mitochondrion-level using high-content imaging analysing ∼1 million mitochondria per condition (Figure 3L). Pharmacological USP30 inhibition results in both an increased number of fused mitochondria (lower width:length) (Figure 3Mi) and a decreased number of fragmented mitochondria (higher width:length) (Figure 3Mii).

These indicate two potential distinct mechanisms by which USP30 inhibitors were able to subtly improve mitochondrial morphology.

### Improvement of mitochondrial morphology by USP30 inhibition occurs independent of PINK1/PRKN-mediated mitophagy

To investigate the dependence of USP30-induced mitochondrial phenotypes on PINK1/Parkin, we generated *PINK1* and *PRKN* knockout SH-SY5Y using CRISPR/Cas9 (Supplemental 2A-B). As expected, loss of PINK1 abrogated depolarization-induced p65Ub formation, whereas loss of PRKN reduced p65Ub by ∼40% as assessed by high-content imaging, western blotting and HTRF-based immunoassay (Figure 4A-B, Supplemental 2C-E). Neither PINK1 KO nor PRKN KO resulted in altered basal mitophagy in SH-SY5Y compared to WT cells as assessed by MitoQC (Figure S2F-G), however, PINK1 KO severely decreased depolarization-induced mitophagy with PRKN KO resulting in a ∼20% decrease in depolarization-induced mitophagy (Figure S2H).

**Figure 4.**
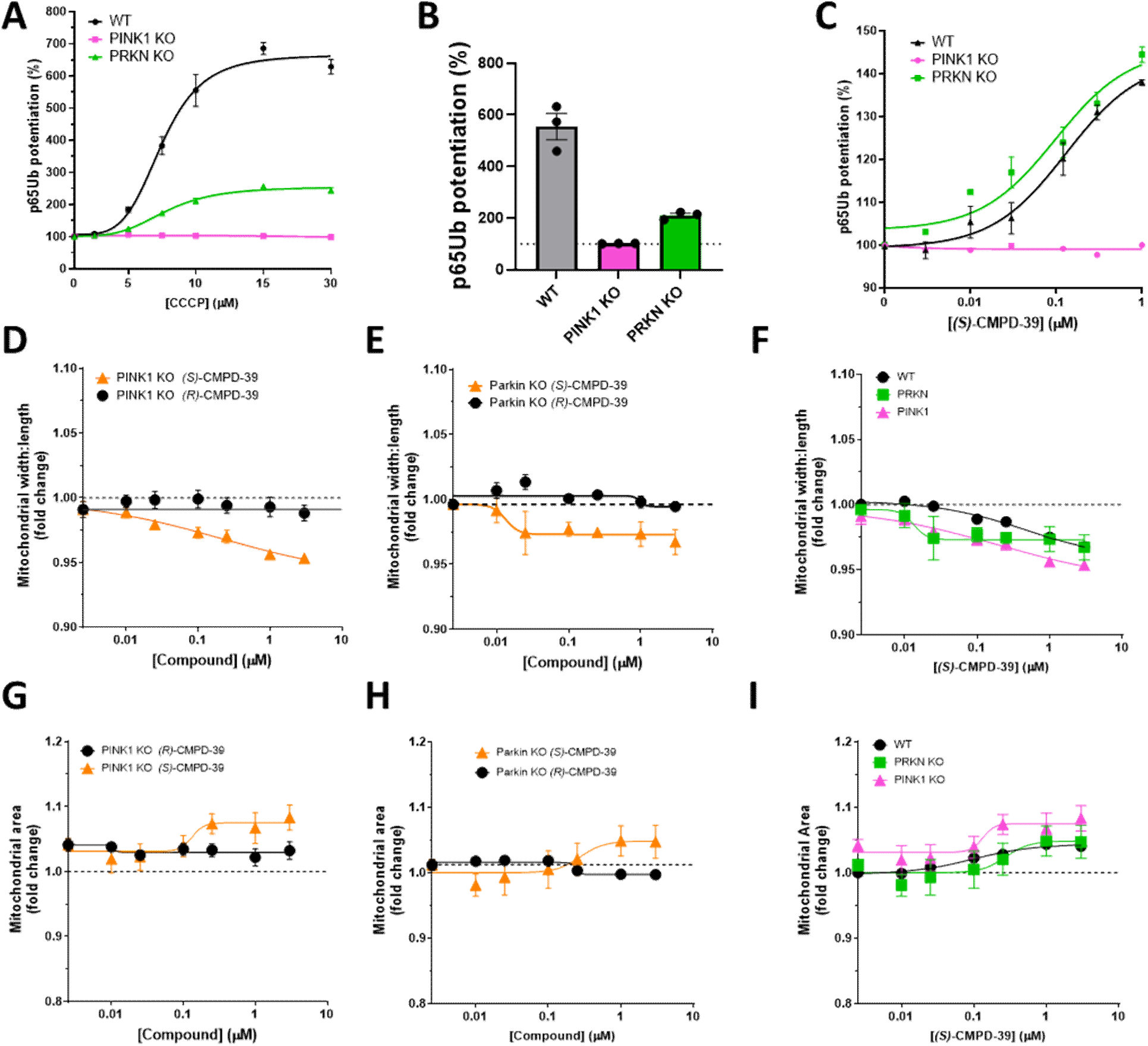
Mitochondrial morphology improvements induced by USP30 inhibition occur independent of PINK1/Parkin. **A-C)** Quantification of p65Ub levels using immunofluorescence in WT, PINK1-/- and PRKN-/- SH-SY5Y lines, normalised to basal p65Ub levels in each line **B)** Quantification of p65Ub potentiation by 10µM CCCP **C)** Potentiation of p65Ub levels by Compound 39 in 10µM CCCP-treated cells normalised within each line. **D-I)** Effect of 48h incubation with *(S)*-CMPD-39 on **D-F)** Mitochondrial morphology and **G-I)** mitochondrial area in WT, PINK1-/- and PRKN-/- lines quantified using MitoQC (n=4-8) **E-F)** Effect of *(S)*-CMPD-39 and compound X in **D/G)** PINK1 KO and **E&H)** Parkin KO lines **F&I)** Comparison of CMPD39 effects in WT, PINK1 KO and PRKN KO lines.

In contrast to PINK1-/-, pharmacological USP30 inhibition resulted in increased p65Ub formation in both WT and PRKN-/- cells. Interestingly, whilst p65Ub generation after CCCP was quantitatively lower in PRKN-/- cells (Figure 4C), the effect of USP30 inhibitors was comparable in magnitude between WT and PRKN-/- cells (Figure 4C), indicating that PINK1 phosphorylation of both Parkin-independent Ub and PRKN-dependent Ub-chains can be potentiated by USP30 inhibition under depolarising conditions.

Given USP30 knockout or pharmacological inhibition alter mitochondrial morphology, we sought to understand if this was dependent on PINK1/Parkin. Knockout of either PINK1 or PRKN did not impair the ability of *(S)*-CMPD-39 to initiate concentration-dependent increases in mitochondrial length (Figure 4D-F) or increases in mitochondrial area in cells (Figure 4G-I) when compared to the effect of *(S)*-CMPD-39 on WT SH-SY5Y, indicating that remodeling of the mitochondrial network is largely independent of the PINK1/PRKN pathway.

### USP30 enhances depolarisation-mediated mitophagy in neuronal models independent of PINK1/PRKN mutation status

To understand the effects of pharmacological inhibition of USP30 in neuronal models we first assessed the effect of *(S)*-CMPD-39 in rat primary cortical cultures. We observed that USP30 inhibition increased mitophagic flux in rat primary cortical cultures, as assessed using mitoQC (Figure 5A-B) consistent with genetic and pharmacological inhibition in SH-SY5Y. Furthermore, p65Ub was assessed in neuronal (MAP2 positive) and glial (GFAP positive) cells within the mixed primary cortical cultures (Figure 5C-D). *(S)*-CMPD-39 potentiated CCCP-induced p65Ub in both neuronal and glial populations with similar EC50∼0.5µM but to a greater extent in neuronal cells (2.2-fold vs 1.5-fold in neurons and glia, respectively).

**Figure 5.**
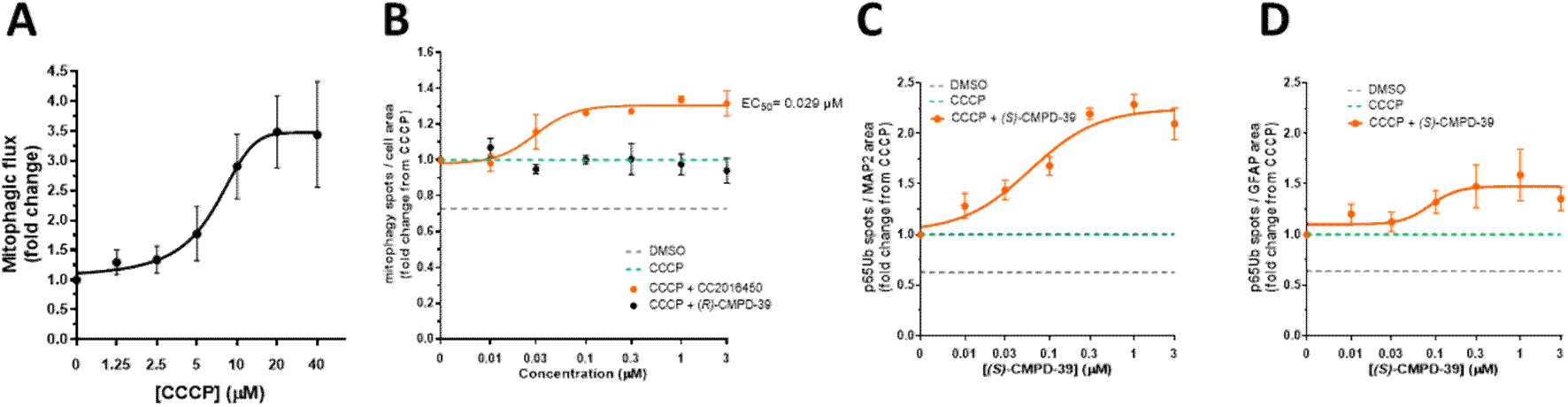
Pharmacological inhibition of USP30 induces mitophagy in primary neuronal cultures. **A)** Induction of mitophagy in primary cortical cultures with CCCP in a concentration-dependent manner (n=4) **B)** Potentiation of CCCP-induced mitophagy by USP30 inhibitor *(S)*-CMPD-39 (n=4). **C-D)** dose-dependent potentiation of p65Ub in **C)** MAP2 positive neurons (n=3) and **D)** GFAP positive cells (n=3).

Given dopaminergic neurons are preferentially vulnerable in Parkinson’s disease, we differentiated iPSC-derived dopaminergic neurons (DaN) using a small molecule approach generating cultures expressing ∼40% midbrain (TH +ve, MAP2 +ve) dopaminergic neurons, as previously described [20, 30] (Figure 6A-B). Treatment of DaN with CCCP (10µM) resulted in a robust upregulation of p65Ub (p<0.001, Paired t-test; n=7;) but was not further potentiated by *(S)*-CMPD-39 (3µM). However, *(S)*-CMPD-39 treatment resulted in a consistent increase in p65Ub levels under basal conditions (Figure S3A-C). These results suggest that under conditions of strong mitochondrial depolarization, USP30 no longer plays a significant role in enhancing mitophagy.

**Figure 6.**
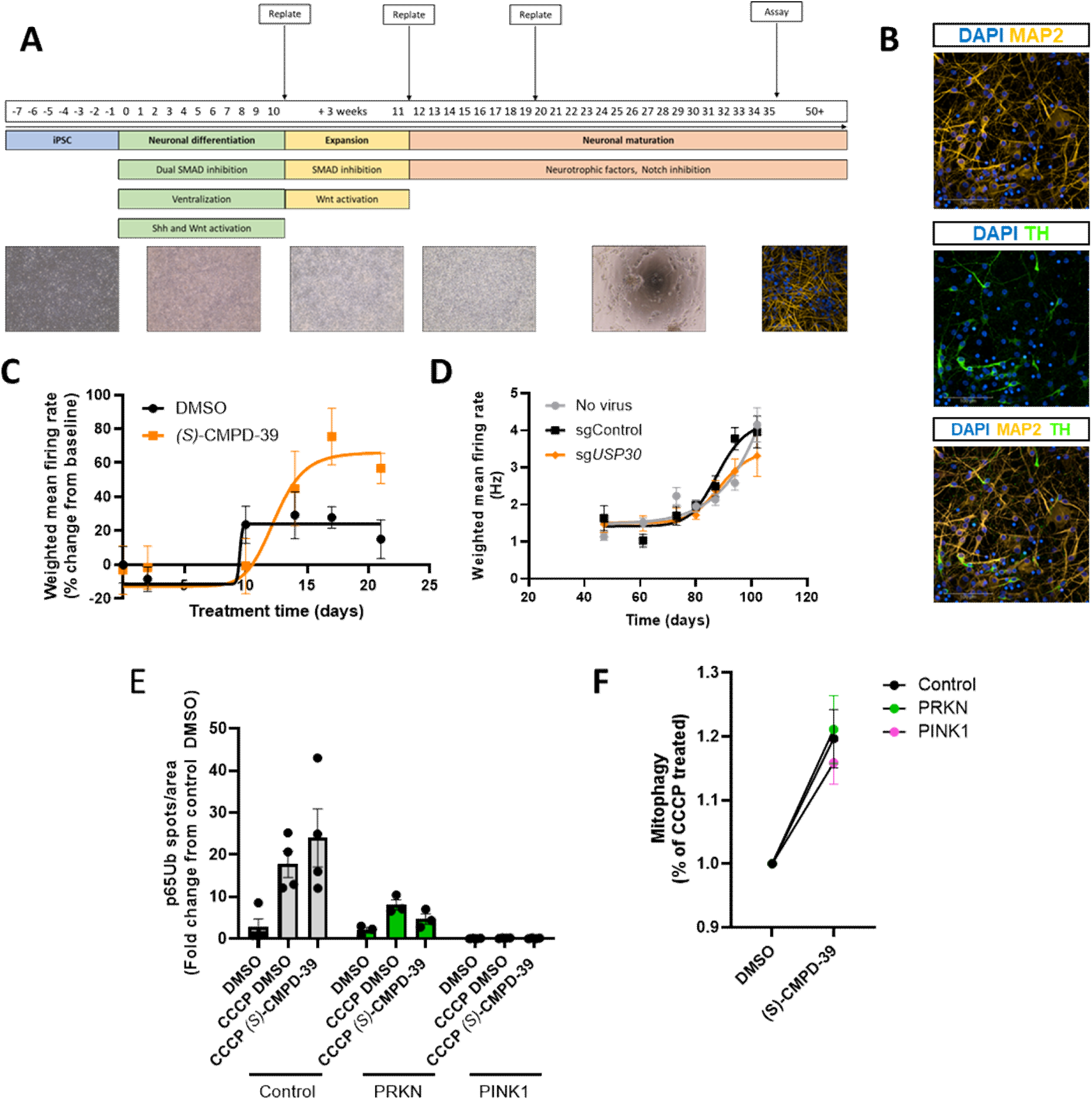
USP30 inhibition potentiates CCCP-induced mitophagy independent of p65Ub induction in iPSC-derived dopaminergic neurons. **A)** Schematic overview of iPSC dopaminergic neuronal differentiation protocol **B)** Representative images of characterisation of iPSC-DaN differentiations staining for MAP2 and tyrosine hydroxylase (TH) co-stained with DAPI. **C-D)** Multi-electrode array analysis of control iPSC-derived neurons **C)** Dopaminergic neurons treated with 3 μM *(S)*-CMPD-39 over 21 days. Weighted mean firing rate (normalised for basal firing rates). **D)** Weighted mean firing rate in dCas9-BFP-KRAB expressing DaN expressing a sgRNA targeting USP30 (sgUSP30) or a non-targeting sgRNA (sgControl) assessed over 55 days from 47-102 days in vitro. E) Effect of USP30 inhibition on p65Ub levels assessed by immunofluorescence in iPSC-derived DaN from control individuals or PD patients with PINK1 or PRKN mutations in the presence/absence of CCCP (6h; 10µM)/*(S)*-CMPD-39 (48h; 3µM). Number of p65Ub spots normalised to control DMSO-treated cells. Potentiation of mitophagy by compound 39 in CCCP-treated iPSC-derived DaN, expressed as a ratio of potentiation from DMSO within genotype.

The electrical activity of dopaminergic neurons has been demonstrated to be closely linked to neuronal health. We therefore investigated the effect of chronic USP30 inhibition on electrical activity in control human iPSC-derived neurons using both pharmacological and CRISPRi approaches. Chronic 21-day treatment of day 82 DaN with 3µM *(S)*-CMPD-39 did not result in large changes to either the firing rate or network activity of the dopaminergic neurons (N=5-6) (Figure 6C & S3D). In line with these results, CRISPRi-mediated knockdown of USP30 did not result in changes in neuronal firing compared to WT or cultures expressing a non-targeting control guide RNA (control), despite USP30 levels being confirmed to be <50% at the end of recording at D107 (Figure 6D, S3E-H). These data suggest that loss of USP30 activity has no overt inhibition of neuronal activity in healthy neurons.

PINK1 and PRKN are well-established genetic causes of Parkinson’s disease and, although biallelic mutations account for a minority of cases overall, they are relatively prevalent in certain populations [36, 37]. To understand if USP30 inhibitors could impact on mitophagy in PINK1 or PRKN patients, DaN were differentiated from iPSCs reprogrammed from Parkinson’s patients with PINK1 or PRKN mutations. As expected, patients with PINK1 mutations demonstrated a lack of production of p65Ub after mitochondrial depolarisation with CCCP (Figure 6E), similarly, PRKN patients demonstrated a decrease in p65Ub production relative to control DaN. USP30 inhibition with *(S)*-CMPD-39 did not rescue p65Ub in PINK1 or PRKN patients either relative to WT cells or when normalised within genotype (Figure S3I). Mitophagy was quantified using mtKeima in control and PINK1 and PRKN patient DaN. Basal mitophagy was not altered in PINK1 or PRKN patient neurons, as assessed using this paradigm and was not significantly enhanced by USP30 inhibition with *(S)*-CMPD-39 (Figure 6F). However, despite line-to-line variation due to differing levels of mtKeima expression, when the effect of *(S)*-CMPD-39 was analysed a on each individual line, *(S)*-CMPD-39 consistently increased mitophagy in CCCP-stimulated DaN independent of PINK1/PRKN mutations (Figure 6F & S3J-k).

Together, these data demonstrate that USP30 enhances depolarisation-induced mitophagy in dopaminergic neurons from both PINK1 and PRKN patients.

## Discussion

Inhibition of the deubiquitylase USP30 is a promising therapeutic strategy for a number of diseases including Parkinson’s. The present study investigates how genetic knockout or pharmacological inhibition of USP30 potentiates mitophagy and affects mitochondrial function in neuroblastoma cell lines, primary neurons and glia as well as human iPSC-derived dopaminergic neuronal cultures.

Decreasing USP30 activity enhanced depolarisation-induced mitophagy in all models. Increases in basal mitophagy were observable with genetic ablation of USP30 and pharmacological inhibition appears to more subtly enhance mitophagy in neuroblastoma and iPSC-derived dopaminergic neuronal cultures. Importantly, pharmacological inhibition of USP30 increased depolarisation-induced mitophagy levels in primary rat neurons and DaN derived from Parkinson’s patients with PINK1 or PRKN mutations comparable to control neurons.

Given the mitophagy-promoting effects of USP30 inhibition, the potential for mitochondrial depletion or deleterious effects of USP30 inhibition has been suggested. Our data in stable USP30^-/-^neuroblastoma cell lines suggests that long-term loss of USP30 activity does not result in mitochondrial impairment but does improve key markers of mitochondrial function consistent with a more efficient mitochondrial network.

Mitochondrial network remodeling was determined by high-content analysis to be a result of subtle changes in both increased mitochondrial network and decreased fragmented mitochondria. These data are consistent with the role of USP30 in increasing the mitophagy of dysfunctional (fragmented) mitochondria and earlier RNAi studies in which mitofusins were demonstrated to play a role in USP30-depletion-mediated mitochondrial fusion [38]. However, in contrast to the shRNA-mediated knockdown of USP30 in HeLa cells which demonstrated a hyper-fused phenotype [38], in the present study, knockout or pharmacological inhibition of USP30 caused subtle changes in mitochondrial networks. In addition, chronic pharmacological inhibition or knockdown with CRISPRi did not significantly affect the firing rate of iPSC-derived dopaminergic neurons.

It has been previously suggested that astrocytes produce greater amounts of p65Ub in response to depolarisation by valinomycin [39]. We observed that USP30 inhibition potentiated p65Ub to a greater extent in neurons compared to GFAP-positive glial cells in rat primary cultures suggesting that specific stimuli may interact with different cell-types to produce varying effects. Furthermore, USP30 inhibition will intersect with this stimulus-cell type to produce varying responses.

These data suggest that USP30 inhibition enhances depolarisation-induced mitophagy largely independently of the rate of p65Ub formation. In addition, the kinetics of mitophagy potentiation by USP30 genetic ablation in neuroblastoma and pharmacological inhibition in iPSC-derived dopaminergic neuronal cultures suggest that the effects of USP30 inhibition are to enhance the earliest stages of mitophagy. This may include upregulation of ubiquitination of substrates around the TOM complex [10] and given the effect of USP30 inhibition was also observed in PINK1 and PRKN patient DaN, it suggests that this process may be reliant on other E3 ligases such as MARCH5 and HUWE1 [10, 40–42]. In the context of previous studies demonstrating that under basal conditions, mitophagy occurs independently of PINK1 in mouse tissues in which there is a high metabolic demand, such as dopaminergic neurons [43], the enhancement of depolarisation-induced mitophagy in PINK1 and PRKN patient lines suggests that priming of mitophagy by USP30 inhibition may still be beneficial in Parkinson’s patients with PINK1 or PRKN mutations. In a similar manner, a recent study has demonstrated that USP30 inhibition induces mitophagy in PRKN KO neurons resulting in decreased oxidative stress in these neurons [17]. In addition, cells lacking PRKN or PINK1 were capable of remodeling mitochondrial networks in response to USP30 inhibition, which is associated with a more efficient mitochondrial network. Our high-content imaging analysis suggests that these effects are at least in part due to decreased numbers of small fragmented mitochondria.

Modest effects of USP30 inhibition in iPSC-derived DaN may be due to near maximal pathway stimulation by CCCP or due to insufficient time (48h) with USP30 inhibition. A previous report in fibroblasts from an individual with a heterozygous Parkin demonstrated that six-day treatment with a USP30 inhibitor was needed to correct mutation-associated p65Ub phenotypes [13] indicating that prolonged inhibition may be needed to observe all phenotypes induced by USP30 knockout.

Our data are in agreement with the trigger-threshold hypothesis put forward by Rusilowicz-Jones and colleagues [44] in which, under basal conditions, USP30 regulates ubiquitylation of OMM substrates located proximally to the TOM complex. These proteins act as ‘seeds’ for ubiquitin chain formation upon mitochondrial depolarisation. Together our data add to the mechanistic rationale for the enhancement of targeted mitochondrial degradation through USP30 inhibition and their potential therapeutic benefit in Parkinson’s disease.

## Acknowledgements

We thank Stewart Humble and Peter Holderieth for producing the sgRNA targeting USP30 and Richard Hargreaves and Mark Labow for their valuable input into the study.

MW, RHR, SNJF, ABM, KS, JC, EB, EM, AMB, KH, WM, PR, AL, HJ and BJR have received salary support from Bristol Myers Squibb through the Oxford DUB Alliance. MB was supported by Michael J Fox Foundation grant SNJF is a recipient of the Oxford-Thatcher Graduate Scholarship. RHR receives the Department of Physiology, Anatomy and Genetics Studentship, the Lady Margaret Hall Alison Brading Scholarship, The Clarendon Fund . This work was supported by the Medical Research Council grant (MR/Y014987/1) and The Rosetrees Trust grant (PhD2024\100031) to BJR.

## Supplemental figures

**Supplemental Figure 1.**
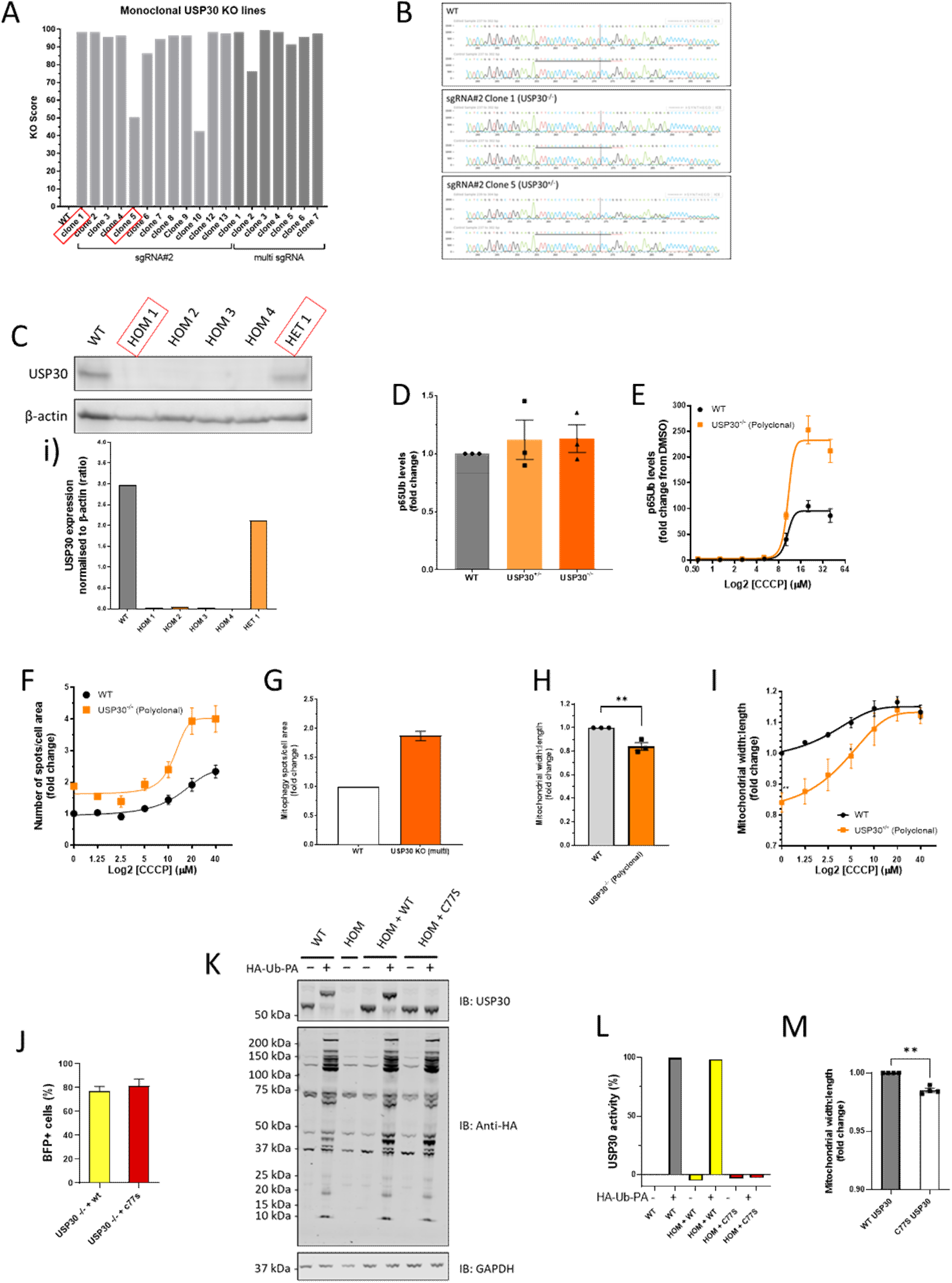
relating to Figure 1. **A)** Inference of CRISPR edits (ICE) score based on sequencing of multiple potential USP30-/- clones **B)** Sanger sequencing traces of WT and identified USP30 heterozygous and homozygous knockout clones. **C)** Western blotting of potential USP30 homo/heterozygous knockout clones **i)** Quantification of USP30 levels in clonal lines. **D)** basal p65Ub levels in USP30^-/-^ and USP30^+/-^ lines as assessed by immunofluorescence (n=3) **E-I)** Effect on mitophagy/mitochondrial morphology of USP30 loss in a polyclonal USP30 KO line **E)** quantification of p65Ub levels in response to CCCP-stimulation (6h) (n=3). **F-G)** Effect of USP30 loss on **F)** basal and **G)** CCCP-stimulated mitophagy as assessed using mitoQC/ **H-I)** Effect of USP30 loss on mitochondrial morphology as assessed using mitoQC under **H)** basal and **I)** CCCP-stimulated conditions. **J)** Quantification of cells expressing transgene in USP30-/-cells infected with USP30 WT or USP30 (C77S) cells. **K-L)** Quantification of USP30 activity using a HA-Ub-PA probe in clonal USP30^-/-^ lines infected with lentivirus encoding WT USP30 or C77S USP30 by western blotting. Activity assessed in lysates by HA-Ub-PA activity assay (n=3-4) **M)** Mitochondrial width:length assessed using mitoQC fluorescence in USP30-/- cells expressing WT USP30 or C77S USP30 (n=4).

**Supplemental Figure 2.**
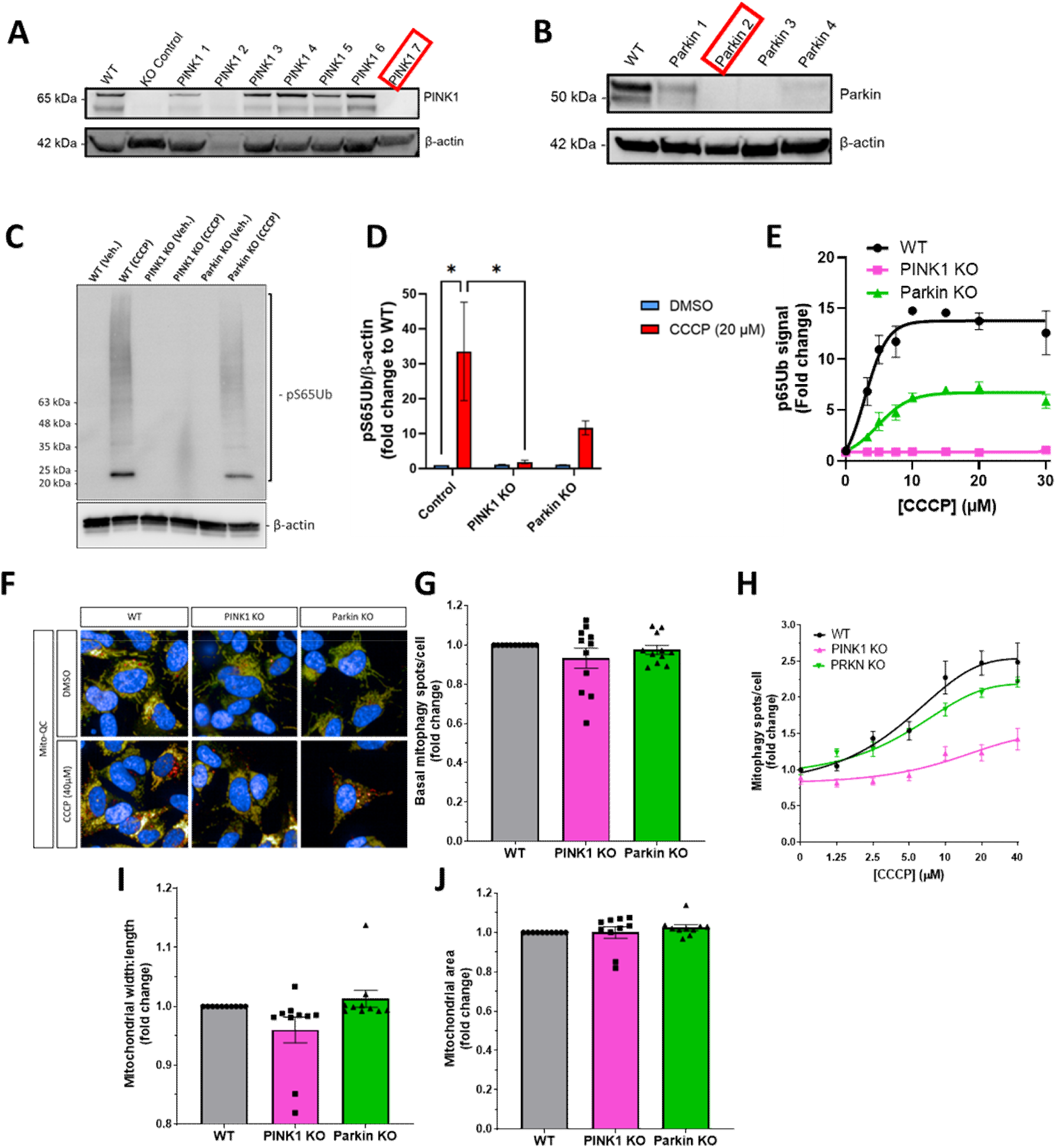
Relating to Figure 4. **A-B)** Western blotting of clonal inducible-Cas9 SH-SY5Y lines derived from polyclonal populations infected with sgRNA lentivirus. **A)** Blotting of candidate PINK1^-/-^ clones **B)** blotting of candidate PRKN^-/-^ clones. **C-E)** Quantification of p65Ub levels by **C-D)** western blot and **E)** HTRF in WT, PINK1^-/-^ and PRKN^-/-^ SH-SY5Y lines, normalised to basal p65Ub levels in WT cells **F-H)** Quantification of mitophagy in WT, PINK1^-/-^ and PRKN^-/-^ lines quantified using MitoQC **F)** Representative images, **G)** basal mitophagy levels (n=11) and **H)** Induction of mitophagy with CCCP (6h) (n=3-6) **I-J)** Effect PINK1/PRKN KO on **I)** mitochondrial morphology and **J)** area (N=10).

**Supplementary Figure 3.**
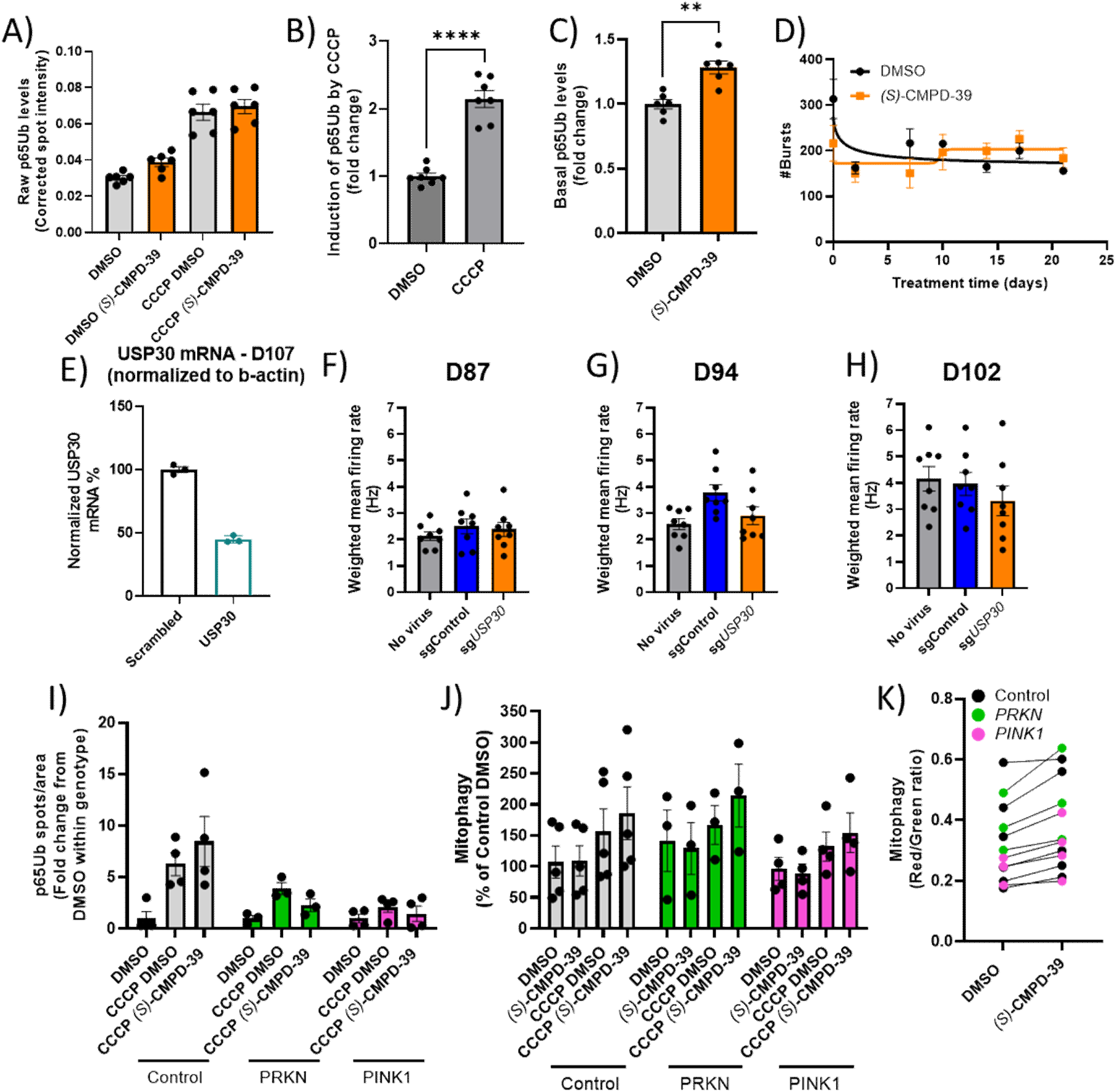
relating to Figure 6. **A-C)** Effect of USP30 inhibition with *(S)*-CMPD-39 (48h (compound replaced after 24h); 3µM) on p65Ub spot intensity levels assessed by immunofluorescence in unstimulated and 10 µM CCCP stimulated iPSC-derived DaN (n=6-7). **B)** Induction of p65Ub by 10 µM CCCP in control iPSC-derived DaN expressed as fold change from average p65Ub spot intensity in DMSO-treated cells. **C)** Effect of USP30 inhibition on raw p65Ub spot intensity levels in unstimulated and 10 µM CCCP-stimulated iPSC-derived DaN from control individuals. **D)** Multi-electrode array analysis of control iPSC-derived neurons treated with 3 μM *(S)*-CMPD-39 over 21 days. Number of network bursts over a 2 min period quantified (n=5). **E)** Quantification of CRISPRi-mediated Knockdown of USP30 in day 107 old neurons (87 days post sgRNA delivery) as assessed by q-RT-PCR, expressed as USP30 expression normalised to ACTB expression normalised to non-targeting sgRNA infected cells (sgControl) (n=3). **F-H)** Weighted mean firing rate (Hz) quantified in non-infected DaN or DaN infected with non-targeting sgRNA (sgControl) or sgUSP30 at **F)** day 87 **G)** day 94 or **H)** day 102 of differentiation. **I)** Effect of USP30 inhibition on p65Ub levels assessed by immunofluorescence in iPSC-derived DaN from control individuals or PD patients with PINK1 or PRKN mutations in the presence/absence of CCCP(6h; 10µM)/Compound 39 (48h; 3µM). Potentiation of p65Ub by CCCP/Compound 39 Number of p65Ub spots normalised within each genotype to DMSO-treated cells**. J)** Effect of USP30 inhibition on mitophagy levels assessed using mtKeima immunofluorescence in iPSC-derived DaN from control individuals or PD patients with PINK1 or PRKN mutations in the presence/absence of CCCP(6h; 10µM)/Compound 39 (48h; 3µM). **K)** Potentiation of mitophagy by Compound 39 in 10µM CCCP-treated cells, as assessed by raw mtKeima pH7:pH4 ratio.

